# Mecp2 deficiency induces dysphagia in a preclinical model of Rett Syndrome

**DOI:** 10.1101/2025.11.19.689305

**Authors:** Luiz Marcelo Oliveira, Maryam Saeed Aslam, Jan Marino Ramirez, Alyssa Huff

## Abstract

Rett Syndrome is a rare, x-linked genetic neurological disorder caused by MECP2 gene mutations. This progressive neurodevelopmental disorder hinders patients’ ability to breathe and eat normally. It is unclear how *Mecp2-*deficiency results in a high percentage of dysphagia and aspiration pneumonia in patients with Rett syndrome. We aim to determine the effects of *Mecp2*-deficiency on swallow related neuromuscular mechanisms contributing to dysphagia in Rett syndrome. Swallow and breathing were detected using electrophysiology in the submental and laryngeal muscle complexes and the hypoglossal and vagus nerves. Several medullary motoneuron populations involved in swallowing were examined by immunohistochemistry in pre and post symptomatic *Mecp2*-deficient male and female mice. Swallow-related submental complex duration and amplitude were significantly decreased in both *Mecp2*-/y and *Mecp2*+/-compared to wild-type, due to decreased motor unit activation. In both *Mecp2-*deficient mice, cholinergic staining in hypoglossal, facial, and trigeminal nuclei were decreased. We noted a significant increase in the transition time from inspiration to swallow, swallow to the subsequent inspiration, and impaired respiratory rhythm regeneration in *Mecp2*-/y, but not *Mecp2*+/- mice.

*Mecp2-*deficiency resulted in impaired brainstem cholinergic signaling, which contribute to weakened submental muscle complex activity, and impaired swallow related laryngeal vestibular closure. These results suggest *Mecp2-*deficient mice are a viable pre-clinical model to further study dysphagia in Rett syndrome.

## Introduction

Whether to pass food to the stomach for nutrition or clear the pharynx for airway protection, swallowing (deglutition) must be done safely to avoid secondary detrimental effects. Swallow is a complex behavior that shares anatomical space with breathing and requires precise temporal synchronization from 26 pairs of muscles across 5 cranial nerves and medullary motoneuron pools to ensure materials enter the esophagus and not the airway (1–3). In order for safe swallowing to occur, the laryngeal elevator muscles (submental complex) must contract to position the epiglottis so it covers the airway, followed by contraction of the laryngeal muscles (laryngeal complex) to then close the airway (4–6). Disruption to laryngeal elevation and closure can result in material penetrating the vocal folds and then aspirating into the lungs. This event can lead to acute infections such as pneumonia—leading cause of death in Rett syndrome (RTT), and chronic maladies like chronic cough, wheezing, respiratory failure, and failure to thrive, which exacerbates underlying conditions and increases the frequency of hospitalizations (7–9).

Difficulty feeding and swallowing have been reported in 81% and 43%, respectively, of patients with RTT. Dysphagia appears to worsen with development, with most patients receiving a diagnosis or intervention after 2 years of age (10–12). Penetration and aspiration of food, liquid, and mucus into the airway, resulting in recurring bouts of aspiration pneumonia was reported in 67-94% of RTT children (11–14). Dysphagia, respiratory distress, pneumonia, malnutrition, growth failure, are major indications of necessary gastrostomy placement (tube feeding) in RTT patients to restore growth, nutrition and health (15). The average age of initial gastrostomy placement averages 8 years of age (15–17), noting a measurable amount of time from symptoms onset to invasive therapeutic intervention, highlighting the need for other therapeutic discoveries.

Fiberoptic endoscopic evaluation of swallow (FEES) along with videofluoroscopic swallow studies (VFSS) are the two gold-standards of clinical testing to detect and determine the severity of dysphagia. The prominent symptoms of dysphagia in RTT are impaired chewing and bolus formation, non-purposeful tongue movement, pharyngeal spillover and ineffective clearance, delayed pharyngeal swallow, and/or delayed laryngeal vestibular closure (11, 12, 18–20). These clinical tests carefully characterize swallow disruption, but provide little insights into the underlying neuromuscular mechanisms. Leading us no closer to understanding how mutation to the methyl-CpG-binding protein 2 (MECP2) gene affects feeding, swallowing and airway protective behaviors.

This study utilizes a preclinical mouse model of RTT-- *Mecp2*^-/y^ knockout (KO) male, *Mecp2*^+/-^ heterozygous (Het) female, and their *Mecp2*^+/y^ and *Mecp2*^+/+^ wild type (WT) counterpart-- to assess the neuromuscular effects of *Mecp2* deficiency on swallow and its coordination with breathing. We hypothesize *Mecp2* deficiency results in altered swallow-breathing timing and impaired laryngeal elevation causing disordered swallow. We aim to identify potential physiologic mechanisms to better understand the interaction between *Mecp2* deficiency and compromised airway protection.

## Methods

### Animals

Heterozygous *Mecp2^+/-^* females (The Jackson Laboratory, Stock No. 003890) were bred with C57BL/6J males (The Jackson Laboratory, Stock No. 000664) to produce both wildtype (*Mecp2*^+/y^ and *Mecp2*^+/+^) and *Mecp2*-deficient (*Mecp2*^-/y^ and *Mecp2*^+/-^) offsprings in which their genotype was determined by PCR analysis. All mice were group housed with *ad libitum* access to food (5053 PicoLab® Rodent Diet, cat no. 3005740-220) and water in a temperature controlled (22 ± 1°C) facility with a 12h light/dark cycle. Both male (WT: p58-p90, KO: p45-p79) and female (WT: p178-p193, Het: p178-p199) mice were used to assess the effects of *Mecp2* deficiency on swallow and its coordination with breathing. A second cohort of mice were used to examine medullary motoneurons in 2-week-old presymptomatic pups. PCR analysis revealed 4 WT, 1 Het, and 3 KO mice. All experiments and animal procedures were approved by Seattle Children’s Research Institute’s Animal Care and Use Committees and were conducted in accordance with the National Institutes of Health guidelines.

### In Vivo Experiments

The same terminal experimental protocol was performed for all 4 groups: ♂WT, ♂KO, ♀WT, ♀Het mice to assess swallow and beathing behaviors using electromyography and electroneurography. However, the age of each experimental group was different due to physiological and experimental factors. For all control mice, the age at experiment was matched to the age of its *Mecp2*-deficient counterpart. Criteria for ♂KO mice experimental date included onset of symptoms such as ataxia, little to no movement in the cage, cold to touch, and/or a body condition score (BCS) of 2 (21). For ♀Het mice, symptoms do not usually appear until 6 months of age, the criteria for experimental date included onset of mild symptoms such as ataxia, and clasping.

As previously published (22), Adult mice were injected with Urethane (1.5 g/kg, i.p. Sigma Aldrich, St. Louis, MO, USA) and secured supine on a custom surgical table. Mice were then allowed to spontaneously breathe 100% O_2_ for the remainder of the surgery and experimental protocol. Adequate depth of anesthesia was determined via heart and breathing rate, as well as lack of toe pinch response every 15 minutes. Core temperature was maintained through a water heating system built into the surgical table. Bipolar electromyograms (EMG) electrodes were placed in the costal diaphragm to monitor respiratory rate and heart rate throughout the experiment. The trachea was exposed through a midline incision and cannulated caudal to the larynx with a curved (180 degree) tracheal tube (PTFE 24 G, Component Supply, Sparta, TN, USA). The recurrent laryngeal nerve (RLN) was carefully dissected away from each side of the trachea before the cannula was tied in and sealed with super glue to ensure no damage to the RLN. A tube filled with 100% O_2_ was attached to the cannulated trachea to provide supplemental oxygen throughout the experiment. The hypoglossal (XII) and vagus (X) nerves were isolated unilaterally, cut distally, and their activity was recorded from a fire-polished pulled borosilicate glass tip (1B150-4, WPI; Sarasota, FL, USA) filled with artificial cerebral spinal fluid (aCSF; in mM: 118 NaCl, 3 KCl, 25 NaHCO_3_, 1 NaH_2_PO_4_, 1 MgCl_2_, 1.5 CaCl_2_, 30 D-glucose) equilibrated with carbogen (95% O_2_, 5% CO_2_), connected to the monopolar suction electrode (A-M Systems, Sequim, WA, USA) and held in a micromanipulator (KITE-R, WPI; Sarasota, FL, USA). Multiple bipolar EMGs, using 0.002” and 0.003” coated stainless steel wires (A-M Systems, Sequim, WA, USA, part no.790600 and 791000, respectively), were simultaneously recorded activity from several swallow and respiratory-related muscle sites.

According to the techniques of Basmajian and Stecko (23), the electrodes were placed using hypodermic needles 30G (part no 305106, BD Precision Glide ™, Franklin Lakes, NJ, USA) in the *submental complex*, which consists of the geniohyoid, mylohyoid and anterior digastric muscles, to determine swallow related laryngeal elevation. The *laryngeal complex,* consisting of the posterior cricoarytenoid, lateral, transverse and oblique arytenoid, cricothyroid and thyroarytenoid muscles, to determine swallow related laryngeal closure. The *costal diaphragm*, used to measure the multifunctional activity for both inspiration, *Schluckatmung* (swallow-breath), a less common diaphragmatic activation during swallow activity (24, 25), and airway protective behaviors such as aspiration reflex (AspR) (Figure 5-figure supplement 1D). Swallow was stimulated by injecting 0.1cc of water into the mouth using a 1.0 cc syringe connected to an oral gavage (22G Instech Labs). At the end of the experiment mice were euthanized by an overdose of anesthetic followed by trans-cardial perfusion (see histology section).

## Analysis

All electroneurogram (ENG) and EMG activity were amplified and band-pass filtered (0.03 – 1 KHz) by a differential AC Amplifier (A-M System model 1700, Sequim, WA, USA), acquired in an A/D converter (CED 1401; Cambridge Electronic Design, Cambridge, UK) and stored using Spike2 software (Cambridge Electronic Design, Cambridge, UK). Using the Spike2 software, data was further processed using a band pass filtered (200-700Hz, 40Hz transition gap) then rectified and smoothed (20ms). Using the Spike2 software, the ECGdelete 02.s2s script is used to remove heart artifact, when present, from the ENG and EMG recordings.

We evaluated swallows that were trigged by injection of water into the oral cavity. *Total swallow duration* was determined by the onset of the submental complex to the termination of the laryngeal complex EMG activity. *Submental complex (SC)* was determined by the onset to the termination of the submental complex EMG activity. *Laryngeal complex (LC)* was determined by the onset to the termination of the laryngeal complex EMG activity. *XII and X duration* was determined by the onset to the termination of each nerve activity. *SC and LC ramp* were determined by the onset to the peak of each muscle activity. *SC and LC* were determined by the peak to the termination of each muscle activity. *Swallow sequence* was calculated as the time difference between the peak activation of the laryngeal and submental complex EMG activity.

*Schluckatmung* (swallow-breath) duration was determined by the onset to the termination of the diaphragm EMG activity during a swallow. *Inspiration-swallow transition* was calculated as the peak of the diaphragm inspiratory activity to the onset of the submental complex. Swallow-*Inspiration transition* was calculated as the offset of the swallow-related laryngeal complex EMG activity to the onset of the subsequent breath. *Diaphragm inter-burst interval (IBI)* was calculated as the offset of the diaphragm EMG activity to the onset of the subsequent breath. All durations are reported in milliseconds (ms) unless otherwise stated. *Respiratory cycle* was calculated as the onset of diaphragm activity during inspiration to the subsequent onset, and measured in seconds (s). Amplitude for SC, LC, diaphragm, XII and X activity was calculated as the % of maximum activity of the muscle/nerve activity of the prior breath (% max of prior breath). For mice that do not have respiratory phasic SC activity, then the maximum activity over the previous 5 respiratory cycles was used.

## Histology

Upon completion of experiments, as previously described (26), animals underwent deep anesthesia using Urethane and were subsequently perfused via the ascending aorta with 20 ml of phosphate buffered saline (PBS 0.1M; pH 7.4), followed by 20 ml of 4% phosphate-buffered paraformaldehyde (0.1 M; pH 7.4), obtained from Electron Microscopy Sciences, Fort Washington, PA. The brains were then extracted and immersed in the perfusion fixative for 4 hours at 4 °C, followed by an additional 8 hours in 30% sucrose solution. Coronal brain sections (30 μm) were obtained using a cryostat and stored in a cryoprotectant solution at -20°C (consisting of 20% glycerol and 30% ethylene glycol and 50% phosphate buffer, pH 7.4) prior to histological processing. All histochemical procedures were conducted on free-floating sections.

For immunohistochemistry (IHC), we first washed three times in PBS 0.1M for 10 min each wash, then Choline acetyltransferase (ChAT) was detected using a polyclonal goat anti-ChAT antibody (AB144P; Millipore; 1:100) diluted in PB containing 2% normal donkey serum (017-000-121, Jackson Immuno Research Laboratories) and 0.3% Triton X-100, incubated for 24 hours. The sections were then rinsed in PBS 0.1M three times for 10 min each, and incubated for 2 hours in Alexa 647 donkey anti-goat antibody (705-605-003; 1:250; Jackson Immuno Research Laboratories. Control experiments confirmed that no labeling was observed when the primary antibodies were omitted. Following staining, the sections were washed in PBS and mounted on slides in sequential rostrocaudal order and allowed to dry in room air for 20 min. The slides were finally covered with Fluoromount-G mounting media with DAPI (00-4959-52; Thermo Fisher) and sealed with a coverslip (150067; Thermo Fisher). For cell counting, imaging, and data analysis, a VS120-S6-W Virtual Slide Scanner (Olympus) was utilized to scan all sections, with images captured using a Nikon DS-Fi3 color camera.

The sections were counted bilaterally, and the numbers reported in the results section correspond exactly to the counts of one-in-two sections of 30-µm sections per mouse, spaced 60 µm apart. The neuroanatomical nomenclature employed during experimentation and section alignment was performed based on the Paxinos and Franklin mouse atlas (27). The number of sections analyzed in each brain region included 5 sections of the dorsal motor nucleus of the vagus (DMV), 7 sections of the XII, 5 sections of postinspiratory complex (PiCo), 7 sections of nucleus ambiguous (NA), 10 sections of the facial (VII) and 6 sections of the trigeminal (Mo5). An image software (Image J, W.S. Rasband/National institute of Health, USA, http://rbs.info.nih.gov/ij) was used for cell counting in 40x magnification photomicrographs. To mitigate potential biases, photomicrography and counting were performed by an anatomist blinded to the experimental conditions.

## Statistical Analysis

A one-way ANOVA followed by Tukey’s multiple comparisons test was used to compare swallow characteristics and its coordination with breathing in all 4 groups (Tables 1 & 4). A two-way ANOVA followed by Tukey’s multiple comparisons test was used to compare histological analysis in all 4 groups (Table 2). Welch’s unpaired t-test was used to compare histological analysis in the pups (Table 3). All data are expressed as mean ± standard deviation (SD), unless otherwise noted. Statistical analyses were performed using GraphPad Prism 10 (GraphPad Software®, Inc. La Jolla, USA). Differences were considered significant at *p* ≤ 0.05.

**Table 1.** Demographics and swallow characteristics in *Mecp2* mice.

|  | WT<br>male<br>mean ( SD )<br>(n=9) | *Mecp2* KO<br>male<br>mean ( SD )<br>(n=9) | WT<br>female<br>mean ( SD )<br>(n=10) | *Mecp2* Het<br>female<br>mean ( SD )<br>(n=9) | p-value |
| --- | --- | --- | --- | --- | --- |
| Weight (g) | 24 ( 2 ) | 14 ( 2 ) | 24 ( 2 ) | 41 ( 10 ) | *male WT vs KO* **0.001**<br>*female WT vs het* **<0.0001**<br>*WT male vs female* >0.99<br>*Mecp2 male vs female* **<0.0001** |
| Age (days) | 74 ( 9 ) | 59 ( 13 ) | 182 ( 4 ) | 189 ( 8 ) | *male WT vs KO* **0.004**<br>*female WT vs het* 0.32<br>*WT male vs female* **<0.0001**<br>*Mecp2 male vs female* **<0.0001** |
| Total Swallow Duration (ms) | 206 ( 57 ) | 161 ( 50 ) | 198 ( 42 ) | 125 ( 19 ) | *male WT vs KO* 0.15<br>*female WT vs het* **0.006**<br>*WT male vs female* 0.98<br>*Mecp2 male vs female* 0.34 |
| Submental Complex Duration (ms) | 168 ( 52 ) | 108 ( 24 ) | 173 ( 39 ) | 99 ( 13 ) | *male WT vs KO* **0.005**<br>*female WT vs het* **0.0004**<br>*WT male vs female* 0.99<br>*Mecp2 male vs female* 0.96 |
| Submental Complex Amp (%max) | 905 ( 453 ) | 351 ( 310 ) | 998 ( 492 ) | 420 ( 260 ) | *male WT vs KO* **0.03**<br>*female WT vs het* **0.02**<br>*WT male vs female* 0.95<br>*Mecp2 male vs female* 0.98 |
| Laryngeal Complex Duration (ms) | 196 ( 54 ) | 157 ( 61 ) | 198 ( 48 ) | 122 ( 21 ) | *male WT vs KO* 0.31<br>*female WT vs het* **0.008**<br>*WT male vs female* 0.99<br>*Mecp2 male vs female* 0.43 |
| Laryngeal Complex Amp (%max) | 781 ( 344 ) | 443 ( 277 ) | 636 ( 243 ) | 391 ( 135 ) | *male WT vs KO* **0.05**<br>*female WT vs het* 0.19<br>*WT male vs female* 0.62<br>*Mecp2 male vs female* 0.97 |
| Hypoglossal Duration (ms) | 206 ( 46 ) | 143 ( 44 ) | 209 ( 49 ) | 122 ( 37 ) | *male WT vs KO* **0.03**<br>*female WT vs het* **0.0009**<br>*WT male vs female* 0.99<br>*Mecp2 male vs female* 0.76 |
| Hypoglossal Amp (%max) | 357 ( 74 ) | 256 ( 184 ) | 369 ( 185 ) | 226 ( 76 ) | *male WT vs KO* 0.46<br>*female WT vs het* 0.14<br>*WT male vs female* 0.99<br>*Mecp2 male vs female* 0.97 |
| Vagus Duration (ms) | 185 ( 48 ) | 137 ( 48 ) | 192 ( 35 ) | 122 ( 20 ) | *male WT vs KO* 0.06<br>*female WT vs het* **0.003**<br>*WT male vs female* 0.98<br>*Mecp2 male vs female* 0.86 |
| Vagus Amp (%max) | 547 ( 294 ) | 406 ( 458 ) | 314 ( 130 ) | 240 ( 107 ) | *male WT vs KO* 0.72<br>*female WT vs het* 0.94<br>*WT male vs female* 0.29<br>*Mecp2 male vs female* 0.60 |
| *Schluckatmung* Duration (ms) | 121 ( 37 ) | 71 ( 15 ) | 125 ( 38 ) | 72 ( 33 ) | *male WT vs KO* 0.14<br>*female WT vs het* 0.30<br>*WT male vs female* 0.99<br>*Mecp2 male vs female* >0.99 |
| *Schluckatmung* Amp (%max) | 53 ( 22 ) | 44 ( 10 ) | 44 ( 7 ) | 86 ( 71 ) | *male WT vs KO* 0.95<br>*female WT vs het* 0.36<br>*WT male vs female* 0.96<br>*Mecp2 male vs female* 0.31 |
| Submental Complex Ramp (ms) | 45 ( 12 ) | 50 ( 15 ) | 45 ( 8 ) | 43 ( 6 ) | *male WT vs KO* 0.71<br>*female WT vs het* 0.97<br>*WT male vs female* >0.99<br>*Mecp2 male vs female* 0.46 |
| Laryngeal Complex Ramp (ms) | 68 ( 33 ) | 71 ( 25 ) | 77 ( 28 ) | 54 ( 11 ) | *male WT vs KO* 0.99<br>*female WT vs het* 0.23<br>*WT male vs female* 0.87<br>*Mecp2 male vs female* 0.51 |
| Submental Complex Decay (ms) | 123 ( 49 ) | 58 ( 17 ) | 128 ( 36 ) | 57 ( 9 ) | *male WT vs KO* **0.0007**<br>*female WT vs het* **0.0002**<br>*WT male vs female* 0.99<br>*Mecp2 male vs female* 0.99 |
| Laryngeal Complex Decay (ms) | 128 ( 38 ) | 85 ( 40 ) | 121 ( 44 ) | 68 ( 16 ) | *male WT vs KO* 0.08<br>*female WT vs het* **0.02**<br>*WT male vs female* 0.98<br>*Mecp2 male vs female* 0.74 |
| Swallow Sequence (ms) | 33 ( 36 ) | 25 ( 13 ) | 32 ( 29 ) | 15 ( 8 ) | *male WT vs KO* 0.89<br>*female WT vs het* 0.41<br>*WT male vs female* 0.99<br>*Mecp2 male vs female* 0.80 |

**Table 2.** Histological analysis of symptomatic *Mecp2* mice.

|  | WT<br>male | *Mecp2* KO<br>male | WT<br>female | *Mecp2* Het<br>female | *p* -value |
| --- | --- | --- | --- | --- | --- |
|  | mean ( SD )<br>(*n*=5) | mean ( SD )<br>(*n*=6) | mean ( SD )<br>(*n*=5) | mean ( SD )<br>(*n*=6) |  |
| Hypoglossal (# ChAT<sup>+</sup> cells) | 460 ( 50 ) | 220 ( 25 ) | 488 ( 22 ) | 234 ( 28 ) | *male WT vs KO* <0.0001<br>*female WT vs het* <0.0001<br>*WT male vs female* 0.63<br>*Mecp2 male vs female* 0.91 |
| Facial (# ChAT<sup>+</sup> cells) | 866 ( 55 ) | 383 ( 80 ) | 860 ( 51 ) | 90 ( 27 ) | *male WT vs KO* <0.0001<br>*female WT vs het* <0.0001<br>*WT male vs female* 0.99<br>*Mecp2 male vs female* <0.0001 |
| Trigeminal (# ChAT<sup>+</sup> cells) | 486 ( 42 ) | 282 ( 27 ) | 500 ( 41 ) | 303 ( 32 ) | *male WT vs KO* <0.0001<br>*female WT vs het* <0.0001<br>*WT male vs female* 0.94<br>*Mecp2 male vs female* 0.75 |
| DMV (# ChAT<sup>+</sup> cells) | 294 ( 32 ) | 301 ( 22 ) | 304 ( 43 ) | 305 ( 26 ) | *male WT vs KO* 0.99<br>*female WT vs het* >0.99<br>*WT male vs female* 0.98<br>*Mecp2 male vs female* 0.99 |
| Nucleus Ambiguus (# ChAT<sup>+</sup> cells) | 230 ( 26 ) | 222 ( 29 ) | 224 ( 22 ) | 221 ( 29 ) | *male WT vs KO* 0.98<br>*female WT vs het* 0.99<br>*WT male vs female* 0.99<br>*Mecp2 male vs female* >0.99 |
| PiCo (# ChAT<sup>+</sup> cells) | 318 ( 33 ) | 308 ( 21 ) | 321 ( 31 ) | 321 ( 39 ) | *male WT vs KO* 0.97<br>*female WT vs het* >0.99<br>*WT male vs female* 0.99<br>*Mecp2 male vs female* 0.93 |

**Table 3.** Histological analysis of pre-symptomatic *Mecp2* mice.

|  | **WT** |  | ***Mecp2* -deficient** |  | ***p* -value** |
| --- | --- | --- | --- | --- | --- |
|  | **mean** | **( SD )** | **mean** | **( SD )** |  |
|  | **(*n* =4)** |  | **(*n* =4)** |  |  |
| Hypoglossal (# ChAT<sup>+</sup> cells) | 472 | ( 28 ) | 414 | ( 81 ) | 0.25 |
| Facial (# ChAT<sup>+</sup> cells) | 803 | ( 82 ) | 560 | ( 179 ) | 0.07 |
| Trigeminal (# ChAT<sup>+</sup> cells) | 489 | ( 44 ) | 439 | ( 37 ) | 0.13 |
| DMV (# ChAT<sup>+</sup> cells) | 252 | ( 52 ) | 290 | ( 31 ) | 0.81 |
| Nucleus Ambiguus (# ChAT<sup>+</sup> cells) | 204 | ( 21 ) | 200 | ( 42 ) | 0.90 |
| PiCo (# ChAT<sup>+</sup> cells) | 325 | ( 29 ) | 301 | ( 19 ) | 0.22 |

Investigators were not blinded during analysis. Sample sizes were chosen based on study logistics rather than determined power analysis.

## Results

### Impaired swallow related neuromuscular transmission

Here we recorded the submental and laryngeal complexes with their corresponding nerves, hypoglossal and vagus, respectively, to assess swallow-related laryngeal elevation and closure in healthy and RTT mice (Figure 1B). A one-way ANOVA with Tukey’s multiple comparisons test identified a significant decrease in submental complex (SC) duration (*p* = 0.005, *p* = 0.0004) and amplitude (*p* = 0.03, *p* = 0.02) in both KO males and Het females compared to WT (Figure 2). While there was a significant decrease in hypoglossal duration (*p* = 0.03, *p* = 0.0009) in KO males and Het females compared to WT, there was no change in amplitude.

**Figure 1:**
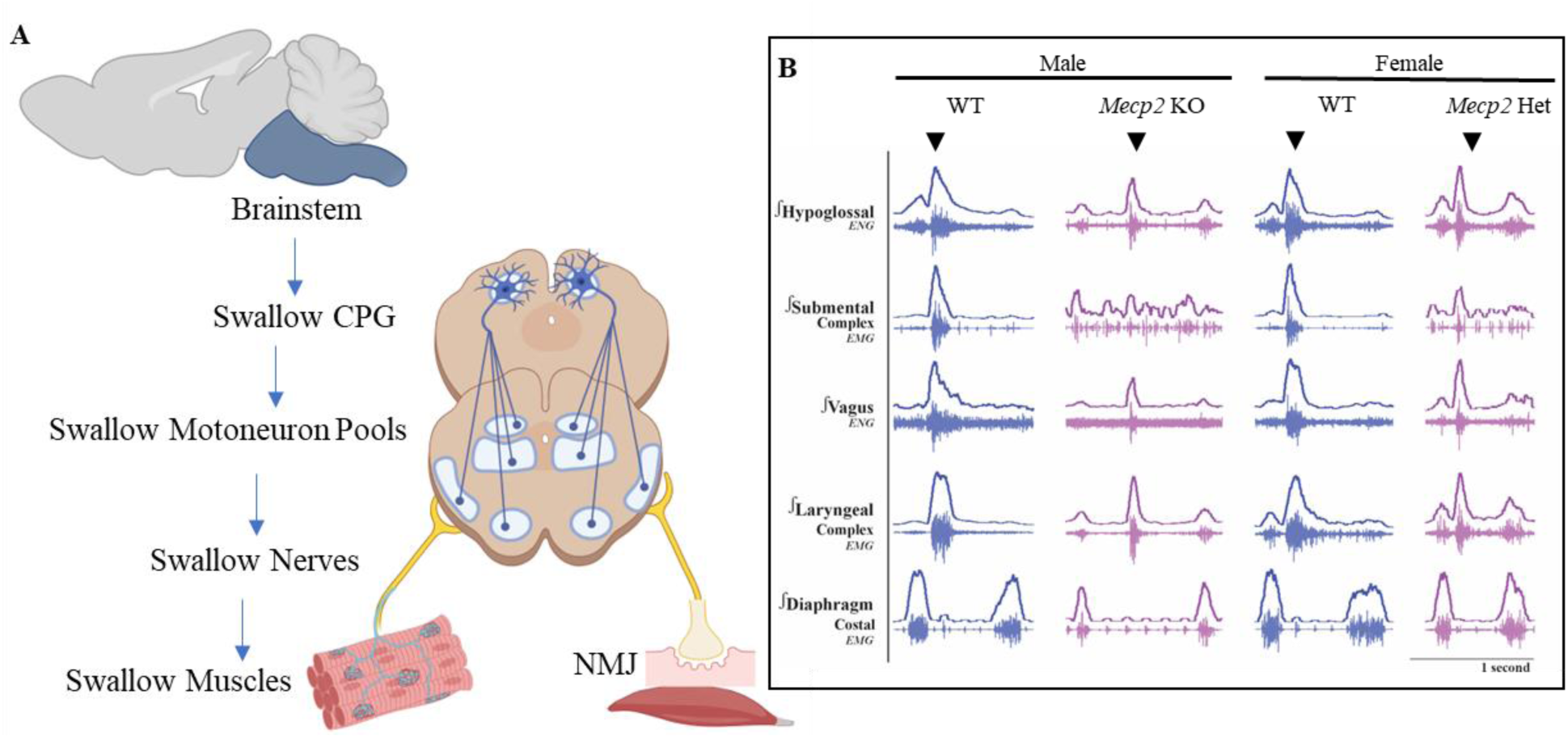
Neuromuscular control of swallow in Rett Syndrome. A) Depiction of motor control of swallow starting with the swallow central pattern generator (CPG) in the medulla to the neuromuscular junction (NMJ) in the muscles. B) Representative traces of swallow (down arrow) and breathing muscles and nerves in all 4 groups of mice: male and female wild type (WT) in blue, male *Mecp2* knock out (KO) and female *Mecp2* heterozygous (Het) in purple. Created in BioRender. Huff, A. (2025) https://BioRender.com/iukaelv

**Figure 2:**
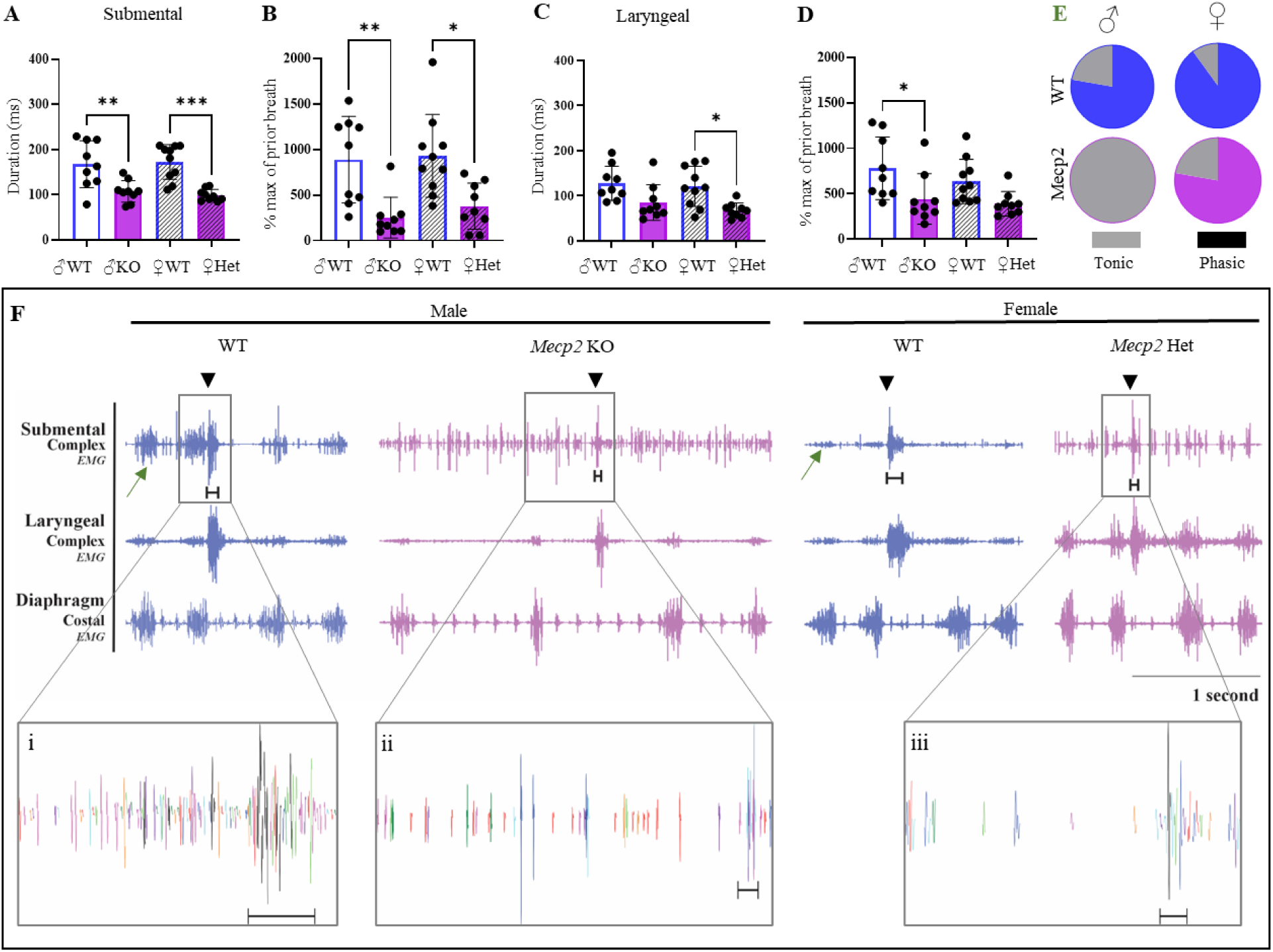
Impaired swallow related submental complex activity due to decrease motor unit activation. Significant decrease in swallow related submental complex (A) duration and (B) amplitude in both KO and Het mice. Significant decrease in swallow related laryngeal complex (C) duration in Het mice and (D) significant decrease in amplitude in KO mice. Respiratory phasic submental complex activity occurred in the majority of WT and Het mice (green arrow in F). While all KO mice had tonic activity, rather than respiratory phasic. F) Representative traces of raw unfiltered spike sorting of submental complex motor units during breathing and swallow showed a decrease in motor unit activity in KO (ii) and Het (iii) mice compared to WT (i). Each color represents a different motor unit.

The decrease in SC duration is due to a significant decrease in the decay duration (*p* = 0.0007, *p* = 0.0002), the time from peak muscle activation to termination, in both KO males and Het females compared to WT (Figure 2- figure supplement 1). While the decrease in amplitude is due to a decrease in number of activated motor units, suggesting impaired or lack of force generation during submental complex activation (Figure 2F). Impaired laryngeal elevation results in impaired swallowing (Figure 3), which could contribute to the significant decrease in body weight (*p* = 0.001) and lifespan (*p* = 0.004) in KO mice (Table 1).

**Figure 3:**
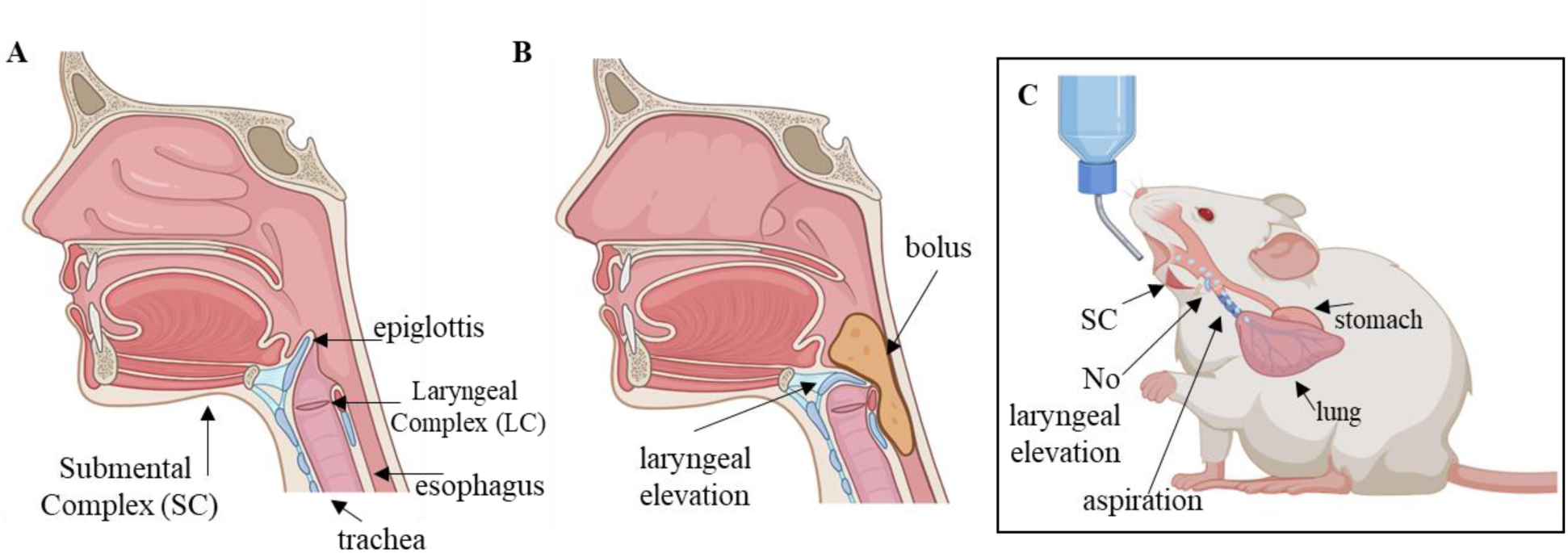
Contraction of submental complex is necessary for laryngeal elevation. The submental complex is a group of laryngeal elevator muscles consisting of the mylohyoid, geniohyoid and digastric. A) At rest the epiglottis, a cartilaginous flap, is positioned away from the larynx to allow for airflow. B) During a swallow the submental complex contracts and the epiglottis is positioned at a downward tilt blocking the airway to prevent the bolus from entering the airway. C) In the event the submental complex is impaired there is limited to no laryngeal elevation and the epiglottis remains in its resting position, and exposes the airway for the bolus to enter and penetrate to the lungs. Panel C is likely what is happening in this study. Created in BioRender. Huff, A. (2025) https://BioRender.com/4r3qqh1

Normally, the SC has respiratory phasic activity, as seen in the Het and WT mice (Figure 2E). However, the SC was not respiratory phasic in the KO mice, rather had tonic activity (Figure 2F).

## *Mecp2* deficiency decreases ChAT expression in swallow related motoneurons

Choline acetyltransferase (ChAT), an enzyme responsible for the synthesize of neurotransmitter acetylcholine (ACh), is used as a maker for motoneurons, due to its’ vital role in neuromuscular transmission (28). Bilateral evaluation of ChAT-expressing neurons detected and assessed swallow-related motoneurons in the brainstem. A two-way ANOVA with Tukey’s multiple comparisons test identified a significant decrease in ChAT^+^ expressing cells in the nucleus of the hypoglossal (*p* < 0.0001, *p* < 0.0001), facial (*p* < 0.0001, *p* < 0.0001), and trigeminal (*p* < 0.0001, *p* < 0.0001) in both symptomatic *Mecp2-*KO and Het mice compared to WT. These motoneurons service the submental complex. However, we saw no change in the number of DMV and NA motoneurons which service the laryngeal complex (Figure 4, Table 2).

**Figure 4:**
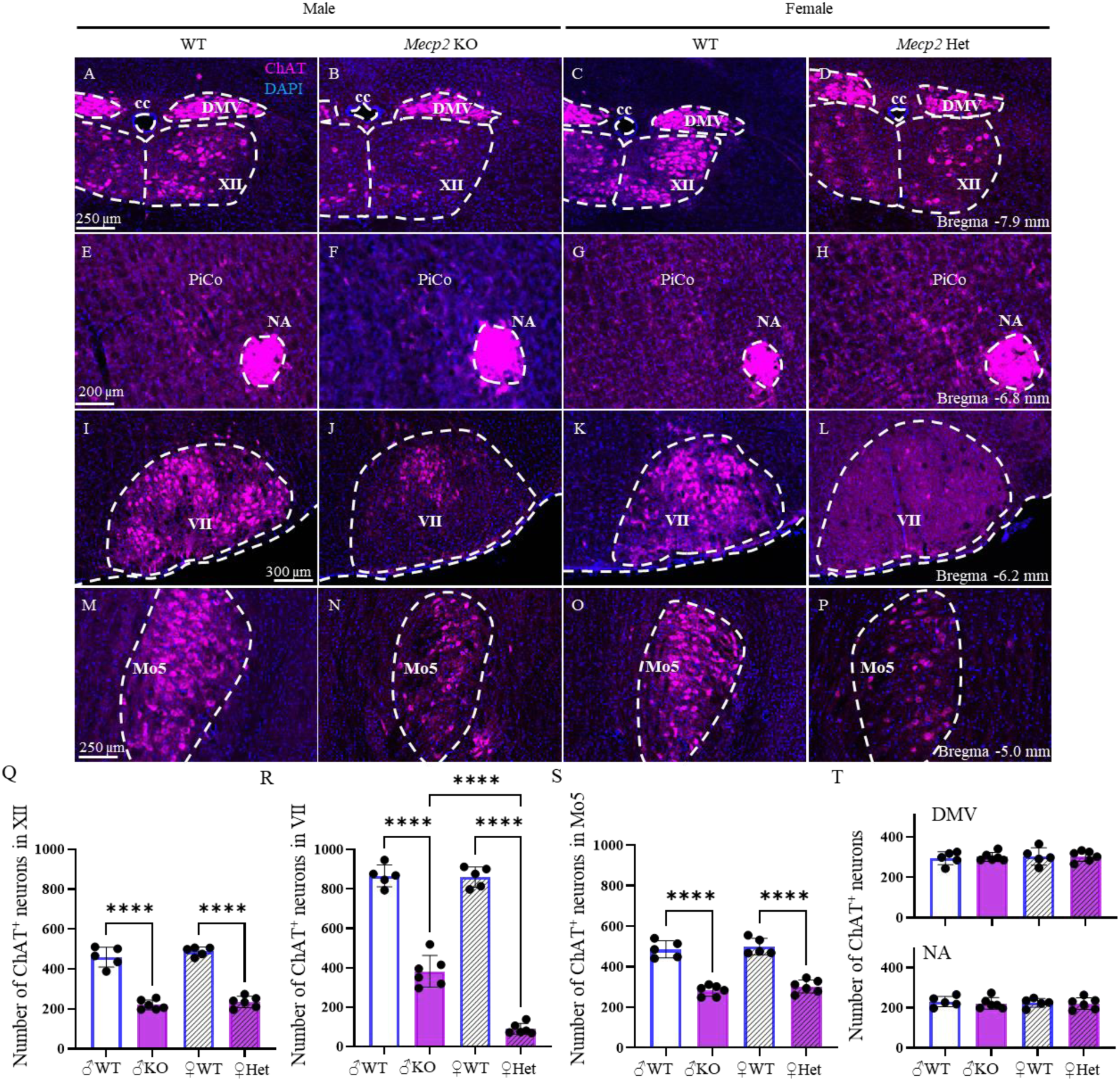
Reduced cholinergic signaling in swallow related medullary motoneurons in symptomatic *Mecp2-*deficient mice. A-P) ChAT (magenta) to detect cholinergic neurons and DAPI (blue) to detect nuclei in the Dorsal motor nucleus of the Vagus (DMV), Hypoglossal motor nucleus (XII), central canal (cc), Postinspiratory Complex (PiCo), Nucleus Ambiguus (NA), Facial motor nucleus (VII), Trigeminal motor nucleus (Mo5). Q) 52% decrease in ChAT^+^ neurons in the hypoglossal nucleus in both KO and Het mice. R) ChAT^+^ neurons were reduced by 56% in KO and 90% in Het in the facial nucleus. S) ChAT^+^ neurons were reduced by 42% in KO and 39% in Het in the trigeminal nucleus. T) No change in ChAT^+^ neurons in the DMV or NA.

**Figure 5:**
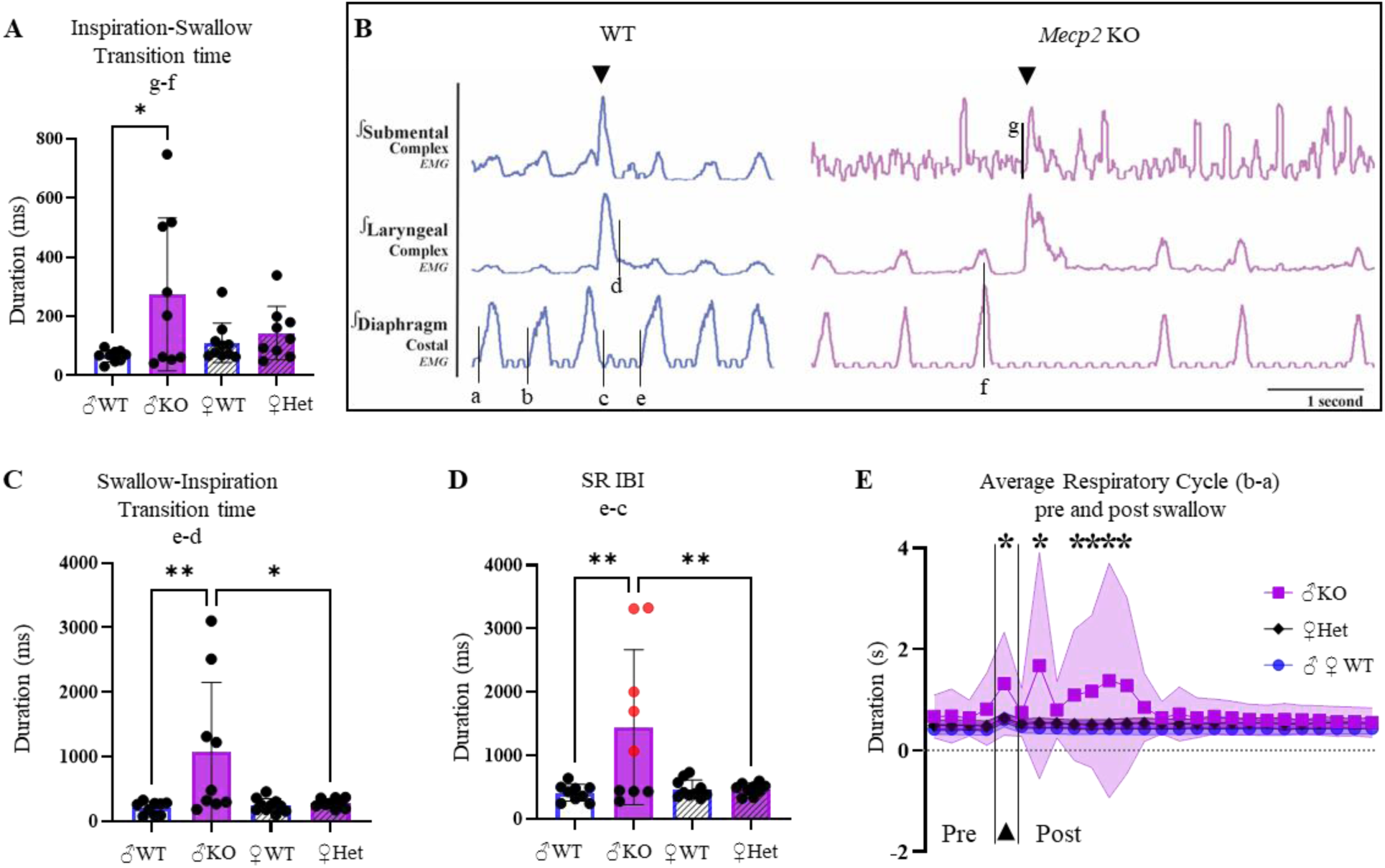
Slowed motor transition between breathing and swallow in *Mecp2* KO, but not Het mice. A) Significant increase in the transition time from inspiration to swallow initiation (g-f) in KO mice. B) Representative traces of swallow and breathing related muscles in WT (blue) and KO (purple) mice showing the respiratory rhythm before and after a swallow. C) Significant increase in the transition time from swallow to inspiration (e-d) in KO mice. D) Significant increase in the swallow related inter burst interval (e-c), also known as the swallow related apnea, in KO mice. Red dots indicate swallows that induced a prolonged apnea defined as quiescence of the diaphragm two times the average duration of the respiratory cycle across 10 respiratory cycles (Figure 5- figure supplement 1). E) Respiratory cycle (RC) duration (b-a) pre and post swallow in male and female WT (blue), Het (black) and KO (purple). There was a significant increase in RC duration during a swallow, however cycles 1 and 3 were of normal duration, but cycles 2, 4-7 were significantly longer, with recovery in most KO mice at cycle 8.

We repeated this in presymptomatic 2-week-old pups and saw no significant difference between WT and *Mecp2*-deficient mice in any of the motoneuron populations (Table 3, Figure 4-figure supplement 1).

## Slowed transition between two neuronal patterns

A one-way ANOVA with Tukey’s multiple comparisons test identified a significant increase in the transition time between inspiration and swallow (*p* = 0.02) as well as swallow and the subsequent inspiration (*p* = 0.008) in *Mecp2* KO compared to WT (Figure 5, Table 4). In five out of nine KO mice, swallow induced apneas (Figure 5D) significantly increased the diaphragm inter-burst interval during a swallow (*p* = 0.006). Apnea was defined as quiescence of the diaphragm two times the average duration of the respiratory cycle across 10 respiratory cycles.

**Table 4.** Swallow-breathing characteristics in *Mecp2* mice.

|  | WT<br>male | *Mecp2* KO<br>male | WT<br>female | *Mecp2* Het<br>female |  | *p* -value |
| --- | --- | --- | --- | --- | --- | --- |
|  | mean ( SD )<br>(*n*=9) | mean ( SD )<br>(*n*=9) | mean ( SD )<br>(*n*=10) | mean ( SD )<br>(*n*=9) |  |  |
| Inspiration-Swallow Transition (ms) | 67 ( 20 ) | 275 ( 258 ) | 111 ( 67 ) | 144 ( 90 ) | *male WT vs KO* | **0.02** |
|  |  |  |  |  | *female WT vs het* | 0.96 |
|  |  |  |  |  | *WT male vs female* | 0.90 |
|  |  |  |  |  | *Mecp2 male vs female* | 0.21 |
| Swallow-Inspiration Transition (ms) | 206 ( 97 ) | 1077 ( 1074 ) | 247 ( 104 ) | 278 ( 72 ) | *male WT vs KO* | **0.008** |
|  |  |  |  |  | *female WT vs het* | 0.99 |
|  |  |  |  |  | *WT male vs female* | 0.99 |
|  |  |  |  |  | *Mecp2 male vs female* | **0.02** |
| SR Diaphragm IBI (ms) | 413 ( 133 ) | 1445 ( 1223 ) | 472 ( 144 ) | 471 ( 92 ) | *male WT vs KO* | **0.006** |
|  |  |  |  |  | *female WT vs het* | >0.99 |
|  |  |  |  |  | *WT male vs female* | 0.99 |
|  |  |  |  |  | *Mecp2 male vs female* | **0.01** |

To evaluate the adaptability of the respiratory network to swallow reconfiguration, we plotted the respiratory cycle duration for four cycles prior, one cycle during, and 21 cycles after swallow. A two-way ANOVA with Tukey’s multiple comparisons test identified no difference in respiratory cycle duration prior to a swallow, rather a significant increase in the 2^nd^, 4^th^-7^th^ cycles (*p* ≤ 0.001) following a swallow in *Mecp2* KO compared to Het and WT mice (Figure 5E). In two out of nine mice, the respiratory rhythm never recovered and death ensued (Figure 5- figure supplement 1). We also saw instances of interrupted laryngeal and diaphragm respiratory synchrony, where the diaphragm missed inspiratory bursts while the laryngeal complex continued (Figure 5- figure supplement 1). Lastly, a putative pharyngeal clearing behavior and a likely aspiration reflex (AspR), characterized as a brief but large burst of diaphragm activity (22), was observed prior to swallowing in two of nine *Mecp2* KO mice (Figure 5- figure supplement 1). None of these events were observed in WT or Het mice.

## Discussion

Deglutition (swallowing) is a brainstem mediated reflex governed by the swallow central pattern generator (CPG) comprised of synchronized bilateral hemi-CPGs (3). Each contains the dorsal swallow group (DSG) located in the caudal nucleus of the solitary tract (cNTS) and the adjacent reticular formation, and ventral swallow group (VSG) located within the ventral reticular formation (29). Multiple mechanisms must properly occur to ensure safe swallowing: 1) The DSG must initiate the swallow command, 2) the VSG then distributes the command to the motor nuclei (2, 30). 3) The motor stimuli must initiate a forceful contraction of appropriate muscles, and 4) the swallow CPG must coordinate with the breathing CPG (25). This study has identified multiple disruptions to these mechanisms in a preclinical mouse model of RTT: 1) reduced cholinergic signaling in swallow motoneurons, 2) impaired neuromuscular transmission in the submental complex, and 3) slowed transition between two motor patterns: swallow and breathing.

There are known cholinergic deficits within the forebrain in girls with RTT (31) and in *Mecp2-*deficient rodents (32, 33). Here we show a cholinergic deficit in the medulla of symptomatic mice, specifically in the hypoglossal, trigeminal, and facial nuclei, responsible for the motor control of the submental complex. However, there were no changes to the DMV and NA in symptomatic mice, as well as no changes to any of the motoneuron populations in presymptomatic mice. This selectivity suggests DMV and NA motoneuron pools may not functionally depend on *Mecp2* expression, and cholinergic signaling is normal prior to symptoms onset.

This study revealed a 52% reduction in ChAT^+^ cells in the hypoglossal nucleus (Figure 4), no change in the amplitude of the hypoglossal nerve (Table 1), and 61% reduction in SC amplitude (Figure 2), which suggest a disruption within the neuromuscular junction (NMJ), likely cholinergic signaling of the N1 receptors (34, 35). The NMJ has been shown to be normal in some muscles of RTT mice (36). Our study suggest that this is not the case for all skeletal muscles as we only see deficits in the submental complex and not the laryngeal complex. Both mouse and human ACh receptors go through a developmental transition between fetal (γ) and adult (ε) subunits (37, 38). Mice lacking the adult subunit have severe muscle weakness and die prematurely (35). It is possible that *Mecp2* plays a role in this transition and contributes to the selective submental muscle weakness in RTT mice.

While this study did not directly measure the force of muscle contraction, rather the activity of the muscle, we are unable to definitively say if the decreased motor unit activation of the submental complex in *Mecp2*-deficient mice resulted in a functional contraction (Figure 2). However, it is reasonable to assume that the decrease in SC amplitude resulted in little to no laryngeal elevation and therefore a lack in laryngeal vestibular closure. In order for laryngeal vestibular closure to occur, the submental muscles must contract, which elevates the larynx and positions the epiglottis over the airway creating a safe environment for the bolus to enter the esophagus and not the trachea (Figure 3B). X-ray swallow evaluations (VFSS) in human RTT patients show impaired laryngeal vestibular closure resulting in supraglottic penetration and glottic aspiration (11). Meaning food/liquid has reached the vocal folds and passed through to airway and into the lungs (Figure 3C) creating an environment prone for developing aspiration pneumonia. While it is unlikely the mice in this study aspirated the water given, due to the placement of a tracheal tube blocking the lower airways, it has been shown that *Mecp2*-deficient mice are capable of aspirating material into the lungs which lead to pulmonary inflammation and damage (39).

PiCo has been shown to play a role in regulating swallow to occur during the postinspiratory phase of breathing (40, 41). While activation of somatostatin (SST^+^) preBötzinger complex (preBötC) neurons can delay the transition between breathing and swallow (25). This suggests alterations to either of these neuronal population could contribute to the transitional delays between swallow and breathing as observed here. However, histological analysis of PiCo does not detect a cholinergic change in *Mecp2-*deficient mice (Table 3), though SST levels positively correlate with *Mecp2* levels in the forebrain (42). Further investigation is necessary to understand SST signaling in *Mecp2-*deficient brainstem.

*Mecp2-*deficient mice and patients have irregular breathing, breath-holds and apneas (43–45). Similar to Voituron et al. (46), we observed central obstructive apneas likely due to a failure of the respiratory CPG to command diaphragmic inspiratory burst, while laryngeal rhythmicity remains (Figure 5- figure supplement 1). Swallow control of the respiratory neural network reconfigures the respiratory network to inhibit inspiration, prolong expiration, and generate swallow apnea (24, 47), but then the respiratory network must seamlessly reconfigure again to generate rhythmic breathing. While we see irregular breathing and apneas during eupnea, integration of swallow into the network further disrupts respiratory rhythmicity, inducing apneas as seen in the RTT patients (48). Most animals are able to regenerate rhythmic breathing after multiple breaths, however for some the network collapses and death occurs (Figure 5- figure supplement 1). Reduction of various neurotransmitters: serotonin (5-HT), gamma-aminobutyric acid (GABA), and norepinephrine (NE); is attributed to variable breathing and apneas in RTT, pharmacological restoration of these systems eliminates breathing abnormalities (43, 49–51).

While the mechanistic understanding of the central reorganization of breathing following a swallow in healthy states has yet to be fully explained, it is likely the slowing of these transitions in *Mecp2-*decifient mice is due to alterations in neurotransmission. The inability of the respiratory network to remain stable during feeding likely contributes to prolonged feeding times, growth failure, apneas/breath-holding (10, 48, 52) seen in RTT children and mice.

Since *Mecp2*^-/y^ testes are internal, they are unable to mate resulting in the inability to generate KO females, *Mecp2*^-/-^. However, Guy et al. created a mouse strain capable of generating KO females and showed that both and female null mice have the same phenotypes (53).

Suggesting that differences seen between KO and Het mice are likely attributed to cellular mosaicism rather than sex differences (54). Here, both KO and Het mice had impairments to SC activity and cholinergic signaling, however only KO mice showed changes in swallow-breathing transitions. KO mice had a 42% reduction in body weight with most having a BCS of 1-2, compared to WT. While Het mice had a 71% increase in body weight with BCS of 5 indicating obesity, compared to WT. Studies evaluating FVB.129F1 *Mecp2*^+/-^ suggest a relationship between hypothalamic *Mecp2* and SST expression and weight gain (42), however this has not been evaluated in C57BL/6J *Mecp2*^+/-^ mice. Hypothalamic *Sim1*-expressing neurons are crucial for energy expenditure and regulating food intake and therefore body weight (55). Deletion of *Mecp2* in these neurons resulted in obesity due to hyperphagia with no change in energy expenditure (56).

There are a few limitations to this study we would like to discuss. We did not normalize neuron number to brain size. However, since we only saw a decrease in ChAT^+^ neurons in a few populations, not all, this decrease is likely not due to a reduction in the size of the brain.

Anesthesia is known to reduce swallow reflex, which resulted in very few sequential swallows in this study. Future studies should evaluate swallow reflex in an awake and alert model to better understand swallow-breathing activity in a more natural feeding session.

Here we have established a preclinical mouse model suitable for studying swallow and airway protection in Rett Syndrome. We have identified a neuromuscular mechanism for impaired laryngeal closure responsible for dysphagia. As well as given insight into possible mechanisms involved in prolonged feeding sessions. Future studies should investigate neuropeptide and neurotransmitter expression in both swallow and breathing medullary centers to better understand swallow control of the respiratory neural network in Rett Syndrome.

## Author Contributions

L.M.O., J.M.R. and A.H designed research; L.M.O., M.S.A., and A.H performed research; L.M.O., M.S.A., and A.H analyzed data; and L.M.O., J.M.R., and A.H wrote the paper.

---

**Figure 2- figure supplement 1:**
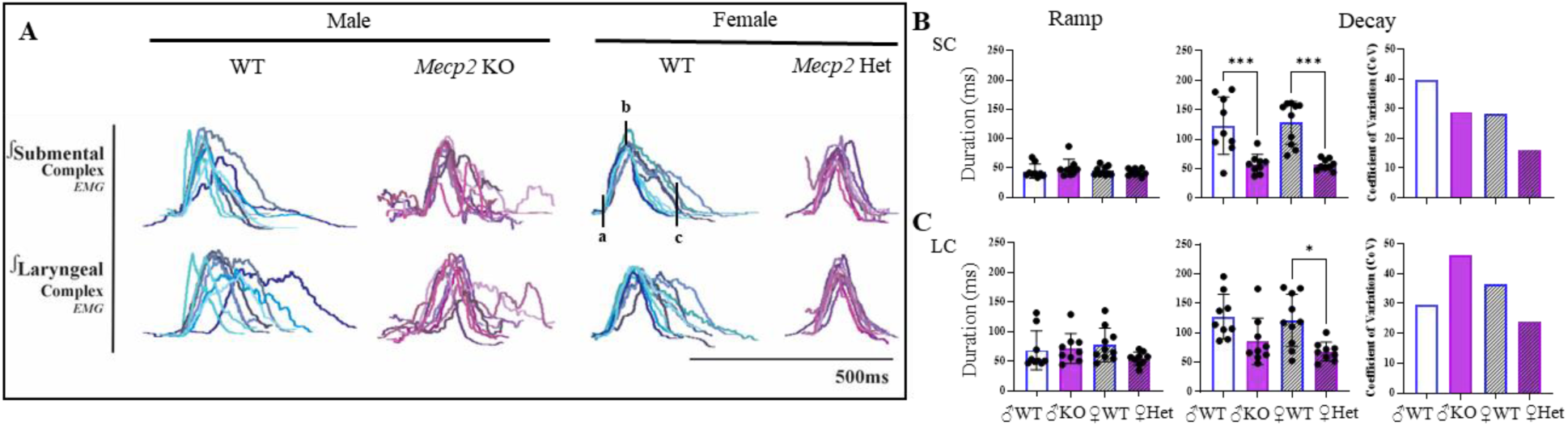
Swallow duration is decreased due to a decrease in submental complex variability and decay duration. A) representative traces of one swallow per animal aligned with the onset of the submental complex to demonstrate the decrease in variability and decay time in the submental complex in WT (blue) and *Mecp2*-deficient (purple) mice. a-b: muscle ramp duration and b-c: decay duration. B) no change in submental complex ramp duration, but a significant decrease in decay duration and variability contributing to the decrease in swallow duration in both KO and Het mice. C) no change in laryngeal complex ramp duration, but a significant decrease in decay duration and variability in Het mice.

**Figure 4- figure supplement 1:**
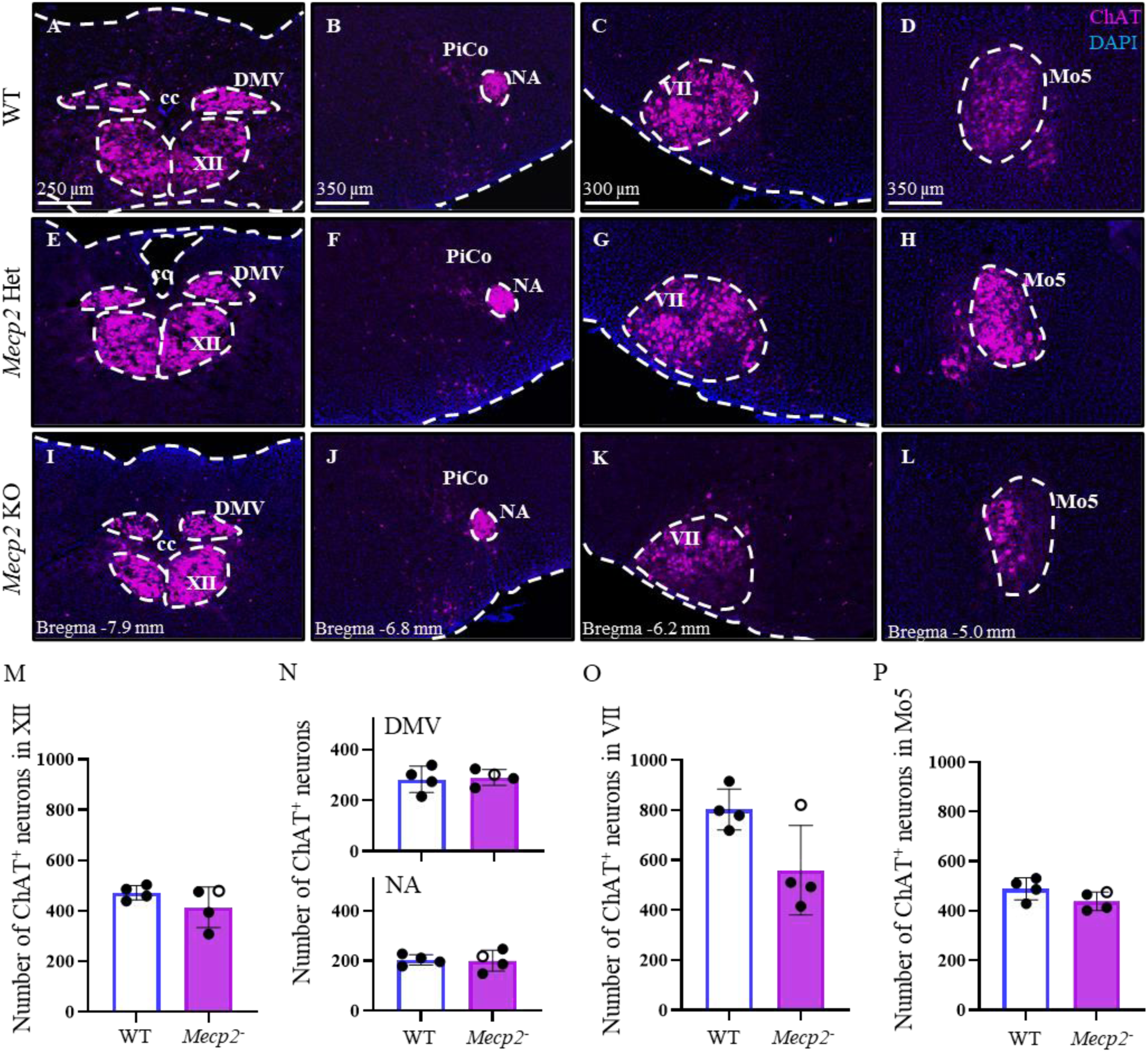
Unchanged cholinergic signaling in swallow related medullary motoneurons in presymptomatic *Mecp2-*deficient mouse pups. A-L) ChAT (magenta) to detect cholinergic neurons and DAPI (blue) to detect nuclei in the Dorsal motor nucleus of the Vagus (DMV), Hypoglossal motor nucleus (XII), central canal (cc), Postinspiratory Complex (PiCo), Nucleus Ambiguus (NA), Facial motor nucleus (VII), Trigeminal motor nucleus (Mo5). Because of the age of the pups, sex was not determined therefor WT (n=4) is likely a mix of male and female. Genotyping indicated n=1 Het female (open circle) and n=3 KO male (closed circle). *Mecp2*-deficient mice were grouped together. We saw no change in ChAT^+^ neurons in M) XII, N) DMV and NA, and P) Mo5. It appears a possible trend in decline of ChAT^+^ neurons in the VII nucleus of KO mice, but not in Het mice, however the low N number prevents from a confident conclusion.

**Figure 5- figure supplement 1:**
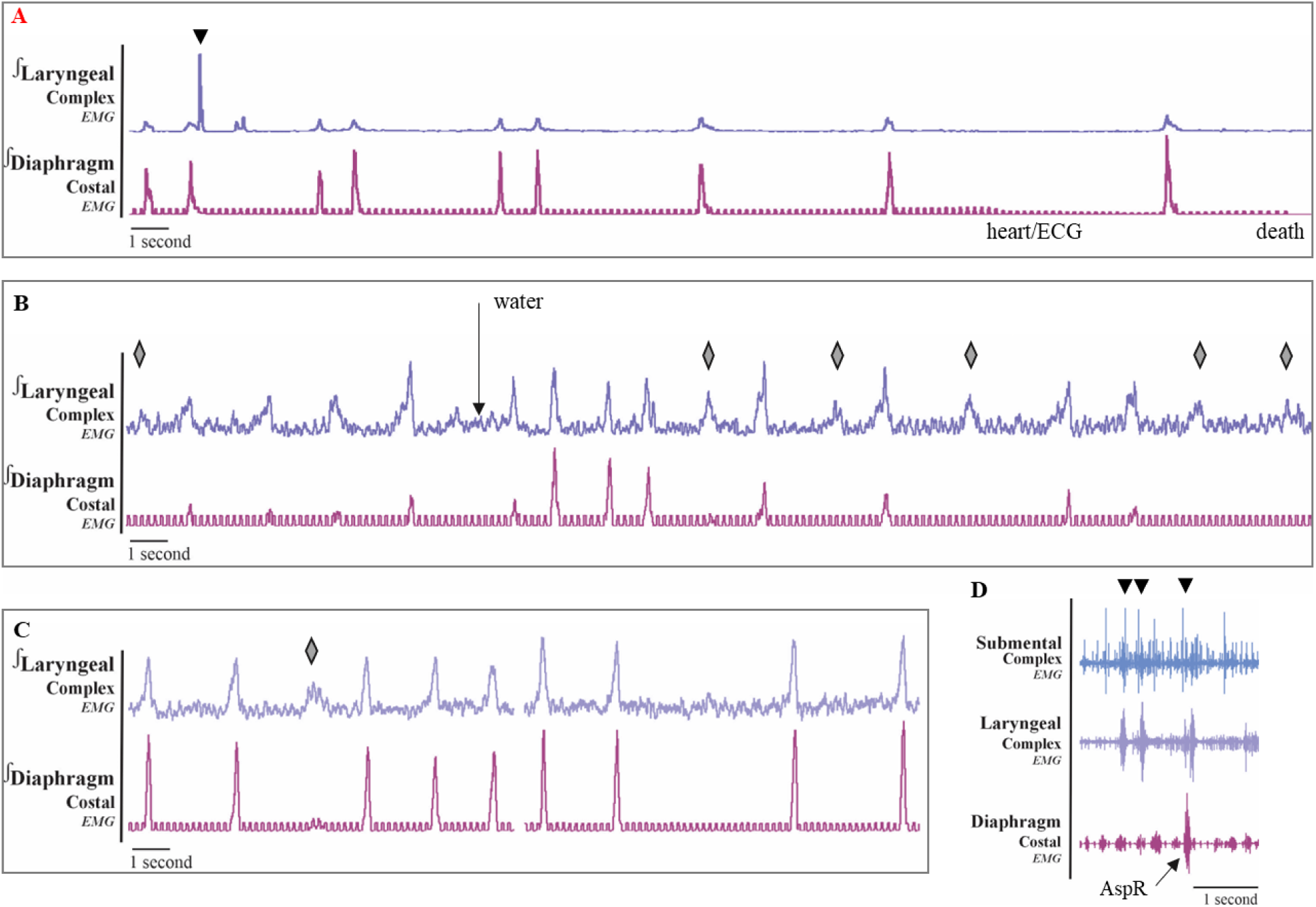
Impaired respiratory neural network reconfiguration leads to death in *Mecp2* KO mice. A) In five out of nine KO mice, swallow induced apneas (Figure 5D). Of those, the respiratory rhythm in two mice was never able to successfully regenerate and resulted in death quickly after the water administration. B) We saw several instances where the diaphragm failed to create an inspiratory burst, while the laryngeal complex rhythmically continued. Due to the amplitude of the laryngeal complex, we do not believe this is a swallow or other laryngeal behavior, rather the bursting activity is inspiration. Since we did not measure airflow, we are unable to determine if airflow was generated during the missed diaphragm bursts. C) We saw these missed diaphragm bursts and apneas in the absence of water stimuli. Diamonds indicate missed diaphragm burst. D) A pharyngeal clearing behavior, likely an aspiration reflex (AspR), characterized as a brief but large burst of diaphragm activity, was seen in two out of nine KO mice. Down arrows indicate swallow.

## References

1. Doty RW, Bosma JF. An electromyographic analysis of reflex deglutition. Journal of neurophysiology 1956; 19: 44–60.

2. Pitts T, Iceman KE. Deglutition and the regulation of the swallow motor pattern. Physiology 2023; 38: 10–24.

3. Jean A. Brain stem control of swallowing: neuronal network and cellular mechanisms. Physiological reviews 2001; 81: 929–969.

4. Burnett TA, Mann EA, Cornell SA, Ludlow CL. Laryngeal elevation achieved by neuromuscular stimulation at rest. Journal of Applied Physiology 2003; 94: 128–134.

5. Vandaele D, Perlman A, Cassell M. Intrinsic fibre architecture and attachments of the human epiglottis and their contributions to the mechanism of deglutition. Journal of anatomy 1995; 186: 1.

6. Sejdić E, Malandraki GA, Coyle JL. Computational deglutition: Signal and image processing methods to understand swallowing and associated disorders. IEEE Signal Process Mag 2019; 36: 138–146.

7. MacKay J, Leonard H, Wong K, Wilson A, Downs J. Respiratory morbidity in Rett syndrome: an observational study. Developmental Medicine & Child Neurology 2018; 60: 951–957.

8. Lotan M, Zysman L. The digestive system and nutritional considerations for individuals with Rett syndrome. The Scientific World Journal 2006; 6: 1737–1749.

9. Ficke B, Rajasurya V, Sanghavi D, Cascella M. Chronic aspiration. 2020.

10. Motil KJ, Caeg E, Barrish JO, Geerts S, Lane JB, Percy AK, Annese F, McNair L, Skinner SA, Lee H-S. Gastrointestinal and nutritional problems occur frequently throughout life in girls and women with Rett syndrome. Journal of pediatric gastroenterology and nutrition 2012; 55: 292–298.

11. Abraham SS, Taragin B, Djukic A. Co-occurrence of dystonic and dyskinetic tongue movements with oral apraxia in post-regression dysphagia in classical rett syndrome years of life 1 through 5. Dysphagia 2015; 30: 128–138.

12. Sideris G, Panagoulis E, Grigoropoulos C, Mermiri D, Nikolopoulos T, Delides A. Fiberoptic Endoscopic Evaluation of Swallowing Findings in Children With Rett Syndrome. Clinical Pediatrics 2023: 00099228231184673.

13. Anderson A, Wong K, Jacoby P, Downs J, Leonard H. Twenty years of surveillance in Rett syndrome: what does this tell us? Orphanet journal of rare diseases 2014; 9: 1–9.

14. Motil KJ, Schultz RJ, Browning K, Trautwein L, Glaze DG. Oropharyngeal dysfunction and gastroesophageal dysmotility are present in girls and women with Rett syndrome. Journal of pediatric gastroenterology and nutrition 1999; 29: 31–37.

15. Motil KJ, Morrissey M, Caeg E, Barrish JO, Glaze DG. Gastrostomy placement improves height and weight gain in girls with Rett syndrome. Journal of pediatric gastroenterology and nutrition 2009; 49: 237–242.

16. Wong K, Downs J, Ellaway C, Baikie G, Ravikumara M, Jacoby P, Christodoulou J, Elliott EJ, Leonard H. Impact of gastrostomy placement on nutritional status, physical health, and parental well-being of females with Rett syndrome: a longitudinal study of an Australian population. The Journal of Pediatrics 2018; 200: 188–195. e181.

17. Berger TD, Fogel Berger C, Gara S, Ben-Zeev B, Weiss B. Nutritional and gastrointestinal manifestations in Rett syndrome: long-term follow-up. European Journal of Pediatrics 2024; 183: 4085–4091.

18. Cocca S, Viviano M, Loglisci M, Parrini S, Monciatti G, Paganelli II, Livi W, Mezzedimi C. Correlation between dysphagia and malocclusion in Rett syndrome: a preliminary study. Sultan Qaboos University Medical Journal 2019; 18: e489.

19. Morton RE, Bonas R, Minford J, Kerr A, Ellis RE. Feeding ability in Rett syndrome. Developmental Medicine & Child Neurology 1997; 39: 331–335.

20. Mezzedimi C, Livi W, De Felice C, Cocca S. Dysphagia in Rett syndrome: a descriptive study. *Annals of Otology*, Rhinology & Laryngology 2017; 126: 640–645.

21. Ullman-Culleré MH, Foltz CJ. Body condition scoring: a rapid and accurate method for assessing health status in mice. Comparative Medicine 1999; 49: 319–323.

22. Huff A, Oliveira LM, Karlen-Amarante M, Ebiala F, Ramirez JM, Kalume F. Ndufs4 inactivation in glutamatergic neurons reveals swallow-breathing discoordination in a mouse model of Leigh Syndrome. Experimental Neurology 2025; 385: 115123.

23. Basmajian JV, Stecko G. A new bipolar electrode for electromyography. Journal of Applied Physiology 1962; 17: 849–849.

24. Pitts T, Poliacek I, Rose MJ, Reed MD, Condrey JA, Tsai H-W, Zhou G, Davenport PW, Bolser DC. Neurons in the dorsomedial medulla contribute to swallow pattern generation: Evidence of inspiratory activity during swallow. PloS one 2018; 13: e0199903.

25. Huff A, Karlen-Amarante M, Pitts T, Ramirez JM. Optogenetic stimulation of pre–Bötzinger complex reveals novel circuit interactions in swallowing–breathing coordination. Proceedings of the National Academy of Sciences 2022; 119: e2121095119.

26. Oliveira LM, Huff A, Wei A, Miranda NC, Wu G, Xu X, Ramirez JM. Afferent and Efferent Connections of the Postinspiratory Complex (PiCo) Revealed by AAV and Monosynaptic Rabies Viral Tracing. J Comp Neurol 2024; 532: e25683.

27. Paxinos G, Franklin KB. Paxinos and Franklin’s the mouse brain in stereotaxic coordinates. Academic press; 2019.

28. Wang W, Salvaterra PM, Loera S, Chiu AY. Brain-derived neurotrophic factor spares choline acetyltransferase mRNA following axotomy of motor neurons in vivo. J Neurosci Res 1997; 47: 134–143.

29. Kessler J, Jean A. Identification of the medullary swallowing regions in the rat. Experimental brain research 1985; 57: 256–263.

30. Bautista TG, Sun Q-J, Pilowsky PM. The generation of pharyngeal phase of swallow and its coordination with breathing: interaction between the swallow and respiratory central pattern generators. Progress in brain research 2014; 212: 253–275.

31. Wenk GL, Hauss-Wegrzyniak B. Altered cholinergic function in the basal forebrain of girls with Rett syndrome. Neuropediatrics 1999; 30: 125–129.

32. Oginsky MF, Cui N, Zhong W, Johnson CM, Jiang C. Alterations in the cholinergic system of brain stem neurons in a mouse model of Rett syndrome. Am J Physiol Cell Physiol 2014; 307: C508–520.

33. Murasawa H, Kobayashi H, Imai J, Nagase T, Soumiya H, Fukumitsu H. Substantial acetylcholine reduction in multiple brain regions of Mecp 2-deficient female rats and associated behavioral abnormalities. PLoS One 2021; 16: e0258830.

34. Carlson AB, Kraus GP. Physiology, cholinergic receptors. 2018.

35. Kalamida D, Poulas K, Avramopoulou V, Fostieri E, Lagoumintzis G, Lazaridis K, Sideri A, Zouridakis M, Tzartos SJ. Muscle and neuronal nicotinic acetylcholine receptors. Structure, function and pathogenicity. Febs j 2007; 274: 3799–3845.

36. Ross PD, Guy J, Selfridge J, Kamal B, Bahey N, Tanner KE, Gillingwater TH, Jones RA, Loughrey CM, McCarroll CS. Exclusive expression of MeCP2 in the nervous system distinguishes between brain and peripheral Rett syndrome-like phenotypes. Human molecular genetics 2016; 25: 4389–4404.

37. Hesselmans L, Jennekens F, Van Den Oord C, Veldman H, Vincent A. Development of innervation of skeletal muscle fibers in man: relation to acetylcholine receptors. The Anatomical Record 1993; 236: 553–562.

38. Cetin H, Beeson D, Vincent A, Webster R. The structure, function, and physiology of the fetal and adult acetylcholine receptor in muscle. Frontiers in molecular neuroscience 2020; 13: 581097.

39. Kida H, Takahashi T, Nakamura Y, Kinoshita T, Hara M, Okamoto M, Okayama S, Nakamura K, Kosai K-i, Taniwaki T. Pathogenesis of lethal aspiration pneumonia in Mecp2-null mouse model for Rett syndrome. Scientific reports 2017; 7: 12032.

40. Huff A, Karlen-Amarante M, Oliveira LM, Ramirez J-M. Role of the postinspiratory complex in regulating swallow–breathing coordination and other laryngeal behaviors. Elife 2023; 12: e86103.

41. Huff AD, Karlen-Amarante M, Oliveira LM, Ramirez J-M. Chronic Intermittent Hypoxia reveals role of the Postinspiratory Complex in the mediation of normal swallow production. Elife 2024; 12: RP92175.

42. Samaco RC, McGraw CM, Ward CS, Sun Y, Neul JL, Zoghbi HY. Female Mecp2+/− mice display robust behavioral deficits on two different genetic backgrounds providing a framework for pre-clinical studies. Human molecular genetics 2013; 22: 96–109.

43. Ramirez J-M, Ward CS, Neul JL. Breathing challenges in Rett syndrome: lessons learned from humans and animal models. Respiratory Physiology & Neurobiology 2013; 189: 280–287.

44. Weese-Mayer DE, Lieske SP, Boothby CM, Kenny AS, Bennett HL, Silvestri JM, Ramirez JM. Autonomic nervous system dysregulation: breathing and heart rate perturbation during wakefulness in young girls with Rett syndrome. Pediatr Res 2006; 60: 443–449.

45. Weese-Mayer DE, Lieske SP, Boothby CM, Kenny AS, Bennett HL, Ramirez JM. Autonomic dysregulation in young girls with Rett Syndrome during nighttime in-home recordings. Pediatr Pulmonol 2008; 43: 1045–1060.

46. Voituron N, Menuet C, Dutschmann M, Hilaire G. Physiological definition of upper airway obstructions in mouse model for Rett syndrome. Respiratory physiology & neurobiology 2010; 173: 146–156.

47. Davenport PW, Bolser DC, Morris KF. Swallow remodeling of respiratory neural networks. Head & neck 2011; 33: S8–S13.

48. Morton RE, Bonas R, Minford J, Tarrant S, Ellis R. Respiration patterns during feeding in Rett syndrome. Developmental Medicine & Child Neurology 1997; 39: 607–613.

49. Abdala AP, Dutschmann M, Bissonnette JM, Paton JF. Correction of respiratory disorders in a mouse model of Rett syndrome. Proceedings of the National Academy of Sciences 2010; 107: 18208–18213.

50. Chen C-Y, Di Lucente J, Lin Y-C, Lien C-C, Rogawski MA, Maezawa I, Jin L-W. Defective GABAergic neurotransmission in the nucleus tractus solitarius in Mecp2-null mice, a model of Rett syndrome. Neurobiology of disease 2018; 109: 25–32.

51. Viemari J-C, Roux J-C, Tryba AK, Saywell V, Burnet H, Peña F, Zanella S, Bévengut M, Barthelemy-Requin M, Herzing LB. Mecp2 deficiency disrupts norepinephrine and respiratory systems in mice. Journal of Neuroscience 2005; 25: 11521–11530.

52. Reilly S, Cass H. Growth and nutrition in Rett syndrome. Disability and rehabilitation 2001; 23: 118–128.

53. Guy J, Hendrich B, Holmes M, Martin JE, Bird A. A mouse Mecp2-null mutation causes neurological symptoms that mimic Rett syndrome. Nature genetics 2001; 27: 322–326.

54. Ribeiro MC, MacDonald JL. Sex differences in Mecp2-mutant Rett syndrome model mice and the impact of cellular mosaicism in phenotype development. Brain research 2020; 1729: 146644.

55. Xi D, Gandhi N, Lai M, Kublaoui BM. Ablation of Sim1 neurons causes obesity through hyperphagia and reduced energy expenditure. PloS one 2012; 7: e36453.

56. Fyffe SL, Neul JL, Samaco RC, Chao H-T, Ben-Shachar S, Moretti P, McGill BE, Goulding EH, Sullivan E, Tecott LH. Deletion of Mecp2 in Sim1-expressing neurons reveals a critical role for MeCP2 in feeding behavior, aggression, and the response to stress. Neuron 2008; 59: 947–958.

